# Deciphering Novel Transcriptional Wiring in Colorectal Cancer: An Integrative Bioinformatic and Experimental Study

**DOI:** 10.64898/2026.08.27.747517

**Authors:** Foad Rommasi, Bahareh Dabirmanesh, Khosro Khajeh

## Abstract

Colorectal cancer remains among the most lethal malignancies worldwide, and the proliferative programme that sustains it has proved to be a challenging target, particularly with acceptable selectivity. Herein, we combined stage-resolved transcriptomic analysis with experimental testing in colorectal cancer cells to inquire whether small molecules, in particular melatonin, act on that programme. The comparison of stage II, III and IV colorectal tumours with normal tissue identified 410 genes upregulated at every stage as a core set, dominated by cell-cycle, spindle-assembly and chromosome-segregation functions. Twenty hub genes were extracted from the corresponding protein interaction network, thirteen of which were required for viability across 59 colorectal cancer cell lines in genome-wide CRISPR screening data. Target-set enrichment nominated E2F4, FOXM1, SIN3A and both DNA-binding subunits of NF-Y as upstream regulators. NF-YA and NF-YB were distinctive in one respect: their annotated targets include *BUB1* and *CCNA2* but exclude *NCAPG*, yielding a testable prediction. Our experimental results showed melatonin reduces SW480 viability with an IC₅₀ of 2.63 mM and lowers *BUB1* and *CCNA2* expression in different manners of concentration-dependency, while *NCAPG* remains unchanged. Melatonin treatment arrests cells in G1 phase, causes a drastic fall in the cycling S-phase fraction, impairs the migration and proliferation phenotype, and rises apoptosis moderately. We also found β2-microglobulin to be an unsuitable normalization reference gene for CRC research due to changes upon treatment. Selective repression of two NF-Y targets with sparing of a non-target is consistent with reduced NF-Y-dependent transcription, though occupancy and subunit-level evidence are to be established.

## 1. Introduction

Cancer is one of the most pressing health challenges of the 21^st^ century. Rather than a single disease, it represents a heterogeneous group of disorders [1] characterized by the uncontrolled proliferation of genetically altered cells. This dysregulated growth disrupts normal tissue architecture and function and can ultimately lead to the spread of malignant cells to distant organs, a process termed metastasis [2]. Colorectal cancer (CRC) has always been ranked among top 5 leading cancers in both genders. At the molecular level, CRC is driven by a range of signalling alterations, genetic mutations, and epigenetic changes, including constitutive activation of the Wnt/β-catenin and RAS–MAPK pathways [3] and TP53 mutations [4]. To support research and clinical management, several classification systems have been developed, notably the Tumour-Node-Metastasis (TNM) system [5] and the Dukes’ staging system [6], which stratify CRC into discrete stages according to the extent of local tissue involvement and distant metastasis.

Melatonin (N-acetyl-5-methoxytryptamine), first isolated in 1958 by Aaron Lerner and colleagues from the bovine pineal gland [7], is an indoleamine with chemical formula of C₁₃H₁₆N₂O₂ and a molecular weight of 232.3 Da [8]. Structurally, the presence of an indole ring confers intrinsic fluorescence to melatonin, characterized by two primary absorption maxima at approximately 224 nm and 290 nm, and a dominant emission peak around 348 nm with shoulder skewed to 450 nm [9]. The electron-dense aromatic system of melatonin facilitates its role as a potent electron donor, enabling efficient neutralization of free radicals and conferring robust antioxidant and ROS-scavenging properties [10].

The pleiotropic actions of melatonin are primarily mediated through engagement with specific G-protein-coupled receptors (GPCRs), comprising two major receptor subtypes: melatonin receptor 1 (MTR1) and melatonin receptor 2 (MTR2), which exhibit distinct tissue distribution patterns and regulatory profiles [11]. Upon receptor binding, melatonin elicits downstream signalling cascades with both receptors coupling primarily to Gi/o proteins. Finally, receptor activation inhibits adenylyl cyclase and lowers intracellular cyclic AMP (cAMP), thereby modulating downstream effectors including protein kinase A (PKA) and the PI3K/Akt pathway [12, 13]. These signals influence transcription for example, through ERK1/2 (extracellular signal-regulated kinase) activity and AP-1 components such as c-Jun, ultimately reshaping gene-expression profiles in target cells [14], which document the effect of melatonin on cell signalling and transcription factors (TFs).

Although recent work has clarified several of melatonin’s mechanisms and signalling pathways, key aspects of its molecular activity, especially those underlying its anti-oncogenic effects, remain incompletely understood. The present study aims to characterize previously undefined molecular mechanisms underlying melatonin’s antitumor effects, using bioinformatic approaches to identify key signalling pathways at the transcriptomic level. We combined differentially expressed gene (DEG) analysis, protein–protein interaction (PPI) network reconstitution, and identification of upstream trans-regulatory factors with *in vitro* experiments in a CRC cell model by including flow cytometry, proliferation assays, and fluorescence-based drug-uptake assays to assess how well the computational predictions agree with experimental data. By integrating these dry-lab and wet-lab approaches, we focused on uncovering novel gene-regulatory effects of melatonin.

## 2. Materials and Methods

### 2.1 Transcriptomic data analysis

A systematic search of the Gene Expression Omnibus (GEO, http://www.ncbi.nlm.nih.gov/geo/) [15] was performed to identify a transcriptomic dataset meeting the study’s inclusion criteria. A microarray dataset from Shunsuke Tsukamoto *et al.* (GEO accession: GSE21510) [16] satisfied the criteria and was selected for downstream analysis as the transcriptomic basis of the study.

To identify DEGs specific to stage II CRC, six laser-capture-microdissection (LCM)-derived tumour samples (patients 046, 057, 063, 082, 107, 139; stage IIA/IIB, Dukes B) served as the experimental group, with matched normal tissue from the same patients as controls. Similarly, for stage III CRC (Dukes C), tumour and matched normal samples from patients 068, 072, 077, 110, and 135 (stage IIIA/IIIB) were analysed. For metastatic CRC (stage IV or Dukes D), the same approach was applied to patients 048, 111, 128, and 134. All differential expression analyses were performed with GEO2R (http://www.ncbi.nlm.nih.gov/geo/geo2r/) [17].

Genes were considered differentially expressed if they met both an adjusted P ≤ 0.05 and |log₂FC| ≥ 2. Subsequent analyses focused on the upregulated subset as druggable overexpressed targets of small molecules, considering their role in promoting aberrant cellular proliferation and tumorigenesis. Next, Venny (https://bioinfogp.cnb.csic.es/tools/venny/) [18] was employed to identify genes consistently upregulated among included stages of CRC.

### 2.2 Network-based identification of hub genes

The PPI network for the shared upregulated genes was built in STRING (Search Tool for the Retrieval of Interacting Genes/Proteins; version 12.0) database (https://string-db.org) [19] using high-confidence interaction score threshold (≥0.7) to ensure the reliability. The resulting PPI network was then imported into Cytoscape software (version 3.10.1; www.cytoscape.org) for network visualization, hub gene identification, and modular analysis. Hub genes, defined as central nodes with high topological importance, were identified by the cytoHubba plugin (version 0.1; https://apps.cytoscape.org/apps/cytohubba) [20] using the Maximal Clique Centrality (MCC) [21], yielding the top 20 hub genes (∼5% of network nodes).

Densely interconnected subnetworks (functional modules) within the PPI network were detected using the Molecular Complex Detection (MCODE) plugin for Cytoscape (version 2.0.0; https://apps.cytoscape.org/apps/mcode). MCODE settings were optimized utilizing the following parameters: Degree Cutoff = 2, Node Score Cutoff = 0.2, K-Core = 2, Maximum Depth = 100, and Haircut post-processing [22]. Integrating cytoHubba and MCODE results, the top 20 genes were retained as the most relevant candidates for CRC progression and potential melatonin-mediated regulation.

KEGG (Kyoto Encyclopaedia of Genes and Genomes) pathway enrichment database (www.genome.jp/kegg) [23] identified pathways associated with the DEGs (adjusted P < 0.05). To identify compounds potentially targeting the shared upregulated genes, the genes were queried against the Proteomics Drug Atlas 2023 from EnrichR (https://maayanlab.cloud/Enrichr/enrich); candidates were ranked by association score, and the top ten drugs (adjusted P < 1.65 × 10⁻²⁵) were shown in a horizontal bar plot.

### 2.3 Bioinformatic analysis of upstream regulatory elements

To characterize the hub genes and identify upstream regulators, including TFs, microRNAs (miRNAs), and metabolites, several resources were used. Functional annotation and enrichment were performed with EnrichR [24], using Gene Ontology (GO) libraries. Regulatory TFs were predicted with ChEA3 (https://maayanlab.cloud/chea3/) using a consensus approach integrating ENCODE and ChIP Enrichment Analysis (ChEA) data (https://maayanlab.cloud/chea3/, https://www.encodeproject.org/software/encode-motifs/) [25]. In parallel, GO EnrichR was utilized to characterize the hub genes in terms of their associated biological processes, molecular functions, and cellular components; thereby, offering insight into their potential roles of hub genes in CRC pathophysiology [24]. Metabolite-gene associations were explored with the Human Metabolome Database (HMDB; https://hmdb.ca/) [26].

The top 20 hub genes were submitted to X2K (Expression2Kinases) database (https://amp.pharm.mssm.edu/X2K/) to infer upstream TFs and kinases [27] and the resulting network was imported into Cytoscape. For interaction modelling, protein/subunit sequences were retrieved from UniProt (https://www.uniprot.org/) [28] and submitted to AlphaFold3 for complex prediction; structures were visualized in Python and UCSF ChimeraX (v1.10.1) [29].

### 2.4 Final verification of bioinformatic workflow

Functional relevance of the hub genes was assessed using gene-essentiality data. The CRISPRGeneDependency.csv file from DepMap Public 22Q4 [30] was obtained from https://depmap.org/portal/download that provides genome-scale CRISPR knockout screening data to assess gene essentiality across a broad panel of human cancer cell lines. Dependencies between the hub genes and CRC cell lines were visualized as clustered heatmaps with CIMminer (https://discover.nci.nih.gov/cimminer/oneMatrix.do) [31]. Prior gene-disease evidence was retrieved from DisGeNET [32].

Prognostic value and expression of the candidate genes were assessed with GEPIA (Gene Expression Profiling Interactive Analysis), which compares tumour and normal tissue using TCGA and GTEx data [33], and with BioGPS, a gene annotation portal providing baseline tissue expression profiles [34, 35], to determine whether these genes are overexpressed in colorectal tumours relative to normal tissue. Finally, immunohistochemistry (IHC) images of normal colon and CRC tissue were obtained from the Human Protein Atlas (HPA) (https://www.proteinatlas.org/) [36] to compare protein expression and localization.

### 2.5 Study hypothesis and experimental design

After identifying the upstream regulators controlling the shared upregulated hub genes, a literature search identified reported interactions between melatonin and these upstream factors [37], linking the computational predictions to potential melatonin-mediated gene regulation in CRC, as an important small molecule.

### 2.6 Cell culture and treatment

The human colorectal cell line SW480 was obtained from the Cell Bank of the Pasteur Institute of Iran (Tehran, Iran). SW480 derives from the primary tumour of a 50-year-old Caucasian male with a stage II colorectal adenocarcinoma. Cells were routinely maintained in high-glucose Dulbecco’s Modified Eagle Medium (DMEM) supplemented with 10% (v/v) fetal bovine serum (FBS; Thermo Fisher Scientific, USA) and incubated at 37 °C in a humidified atmosphere containing 5% CO₂.

Melatonin powder was kindly provided by Jalinous Pharmaceutical Co. (Tehran, Iran). A concentrated stock solution of melatonin (100mM) was prepared by dissolving the compound in pure ethanol (Merck Millipore, USA), followed by serial dilution in serum-free DMEM to achieve the desired working concentrations. Cells were cultured in serum-free media during the treatment to avoid potential interactions of serum proteins with melatonin over the course of treatment.

### 2.7 Cell viability assay

Melatonin cytotoxicity was assessed by MTT assay. Six concentrations (20, 10, 5, 2.5, 1.25, 0.63 mM; twofold serial dilution) and vehicle control were tested [38]. Briefly, 10,000 SW480 cells were seeded into each well of a flat-bottom 96-well microplate (SPL Life Sciences, South Korea). After cell attachment, the culture medium was replaced with 200 μL of culture media containing the respective melatonin concentrations (no FBS). Following 48 hours of treatment, the medium was replaced with 100 μL of MTT solution (5 mg/mL), prepared by dissolving methylthiazolyldiphenyl-tetrazolium bromide in phosphate-buffered saline (PBS), and cells were incubated 4 h to form formazan.

After incubation, the resulting formazan crystals were solubilized by adding dimethyl sulfoxide (DMSO), and absorbance was measured at 570 nm (formazan absorption peak) and 630 nm (background correction for non-specific absorbance by cellular debris or aggregates) using a BioTek microplate reader (Agilent Technologies, USA). Cell viability was calculated using a normalization approach **(Supplementary 1A)**, and solvent controls received ethanol at the same v/v ratio used for melatonin treatment. From the dose–response fit (GraphPad Prism v9.5.1; **Supplementary 1B**), two concentrations were selected: IC₆₅ (65% viability loss; 35% viable) and IC₃₅ (35% viability loss; 65% viable).

### 2.8 Real-time PCR

For transcriptional profiling after melatonin treatment, total RNA was extracted from SW480 cells treated with IC₃₅ or IC₆₅ and vehicle untreated group. For each condition, 1.5 × 10⁶ cells were seeded in T25 flasks (SPL Life Sciences, South Korea) at ∼50% confluency, matching the MTT conditions. After 48 h, RNA was isolated with TRI Reagent (Sigma-Aldrich, USA) per the manufacturer’s protocol [39], resuspended in nuclease-free DEPC-treated water and assessed by 2% agarose gel electrophoresis and NanoDrop ND-2000 spectrophotometry (Thermo Fisher Scientific, US). Complementary DNA (cDNA) synthesis was performed using high-quality RNA templates, BioFact™ RT Kit reagents, and random hexamer primers (BioFACT, South Korea), as described elsewhere [40]. Gene-specific primers for both target and reference genes were designed using OligoAnalyzer (https://eu.idtdna.com/pages), and further evaluated using NCBI Primer-BLAST (https://www.ncbi.nlm.nih.gov/tools/primer-blast/) to ensure specificity and efficiency [41]. The sequences of primers used in this research are provided in **Table 1**.

**Table 1.**
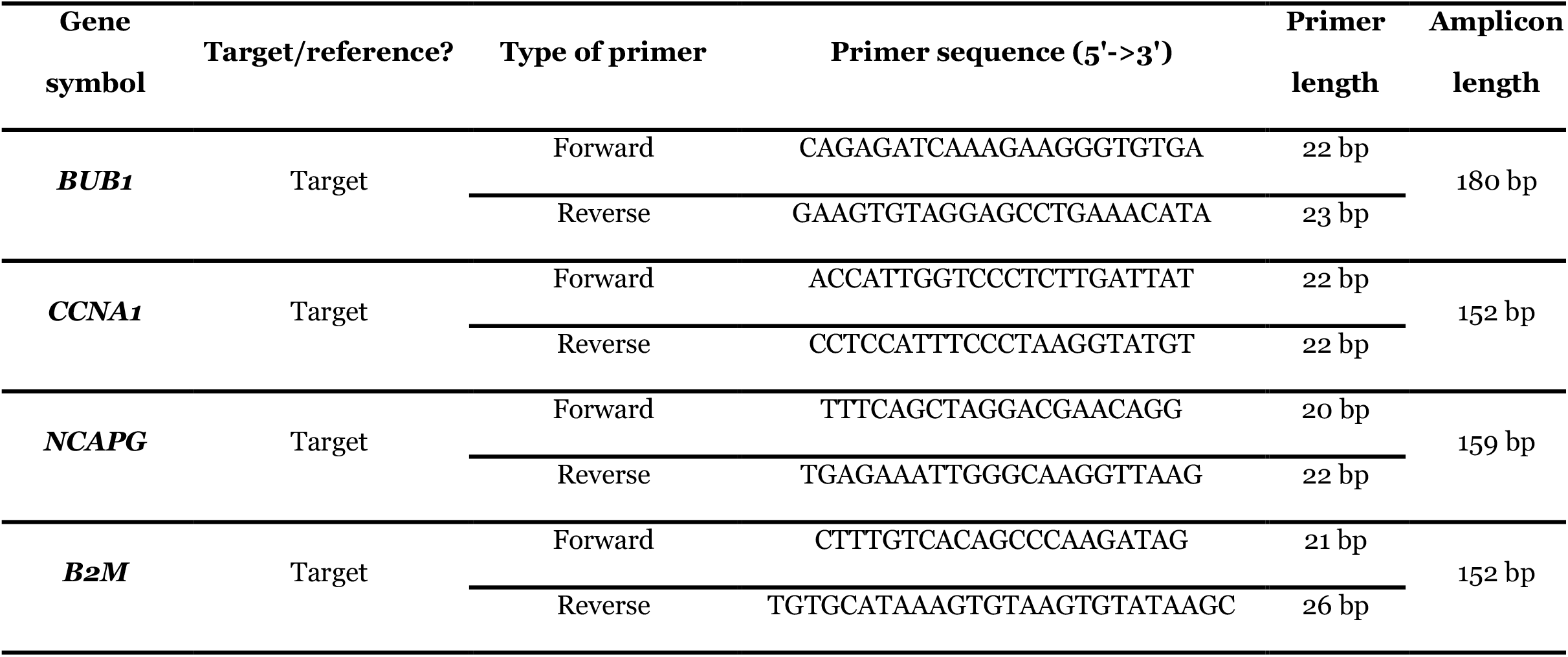
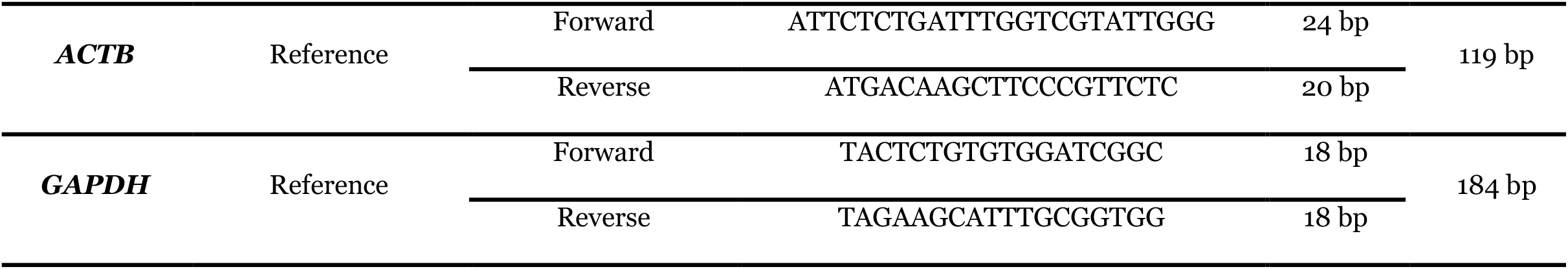
The primers designed for the candidate genes expression analysis by Real-time PCR.

### 2.9 Quantitative gene expression analysis

Quantitative real-time polymerase chain reaction was employed to measure transcriptional changes in candidate genes following the treatment, as described previously with slight modifications [42]. Quantification of relative gene expression was conducted using the threshold cycle (Ct), normalized against the geometric mean of two internal reference genes (GAPDH and β-actin), which were selected based on their consistent expression in colorectal cancer cell lines [43]. Relative fold changes were calculated using the Pfaffl method [44], which accounts for gene-specific amplification efficiencies as presented in **Supplementary 1C**.

### 2.10 Cell proliferation and migration assay

Effects of melatonin on SW480 migration were assessed by wound-healing (scratch) assay [45]. Briefly, 3 × 10⁵ SW480 cells were seeded into each well of a 12-well plate (SPL Life Sciences, South Korea) and incubated for 48 hours to reach approximately 90% confluency. Afterwards, a linear scratch was made, and detached cells were rinsed away with PBS. Treated groups received complete medium with IC₃₅ or IC₆₅; untreated received medium with the vehicle. Images were captured at 0, 24, and 48 h (phase-contrast inverted microscope; ZEISS, Germany), and wound closure was quantified in ImageJ (version 1.52v, NIH, US).

### 2.11 Fluorescence-based internalization assay

To test how melatonin influences intercellular fluorescence, the internalization assay was done [46]. Shortly, 5.2 × 10⁵ SW480 cells, calculated to achieve confluency comparable to that used in the MTT assay, were seeded into 6-well black-walled plates with optically clear, crystal quartz bottoms and incubated overnight. The following day, test groups were treated with previously defined IC₃₅ and IC₆₅ concentrations of melatonin, while untreated groups received the vehicle. After 48 hours of incubation, culture media were aspirated and the cells were washed twice with PBS to eliminate extracellular melatonin and reduce background fluorescence.

Imaging used a confocal laser scanning microscope (Olympus, Japan), exploiting melatonin’s intrinsic fluorescence (excitation 405 nm, emission 455 nm). Intensity was quantified in ImageJ software. Corrected Total Cell Fluorescence (CTCF) was calculated using the equation in **Supplementary 1D**, correcting the background fluorescence and cell area, enabling quantitative comparison of melatonin internalization across groups.

### 2.12 Flow cytometry analyses

Herein, flow cytometry was utilized to complement gene expression analysis by evaluating the functional consequences of melatonin treatment in SW480 cells in terms of apoptosis induction, cell cycle arrest, and oxidative stress response [47]. Cells were treated using the same protocol as described for RNA extraction. Following a 48-hour incubation, cells were detached using Accutase (Capricorn Scientific, Germany) to preserve membrane integrity and surface markers. Subsequent procedures, including staining protocols for apoptosis detection (Annexin V/PI), cell cycle distribution (propidium iodide staining for DNA content), and intracellular ROS quantification (DCFH-DA assay), were conducted in accordance with standardized flow cytometry protocols, as previously described elsewhere [48].

### 2.13 Statistical analysis and data visualization

All experiments used at least two independent biological replicates, each with at least three technical replicates. Data are presented as mean ± SD. Analyses were performed in GraphPad Prism v9. Two groups were compared by two-tailed Student’s t-test; multiple groups by one-way ANOVA with Tukey post hoc correction. *P* (or adjusted *P*) < 0.05 was considered significant. Visualization used GraphPad Prism, MS Office, and ImageJ.

## 3. Results

### 3.1 Transcriptomic profiling and network-based prioritization identify candidate drivers of CRC progression

Comparative analysis of DEGs in stage II, III and IV CRC relative to normal colorectal tissue identified 929, 1113 and 691 upregulated genes, respectively **(Table 2)**.

**Table 2.** Number of DEGs in various stages of CRC.

| Group number | Stage | DEG type | Number of DEGs |
| --- | --- | --- | --- |
| Group 1 | Stage 2 CRC | Upregulated | 929 |
|  |  | Downregulated | 745 |
| Group 2 | Stage 3 CRC | Upregulated | 1113 |
|  |  | Downregulated | 813 |
| Group 3 | Stage 4 CRC | Upregulated | 691 |
|  |  | Downregulated | 267 |
| The number of shared upregulated genes among Group 1, 2 and 3 |  |  | 410 |
| The number of shared downregulated genes among Group 1, 2 and 3 |  |  | 195 |

Of the 1,463 genes upregulated in at least one stage, 410 (28.0%) were upregulated in all three, defining a core transcriptional programme sustained throughout disease progression, whereas 142 (9.7%), 322 (22.0%) and 139 (9.5%) genes were upregulated exclusively in stage II, III and IV, respectively **(Figure 1A)**.

**Figure 1.**
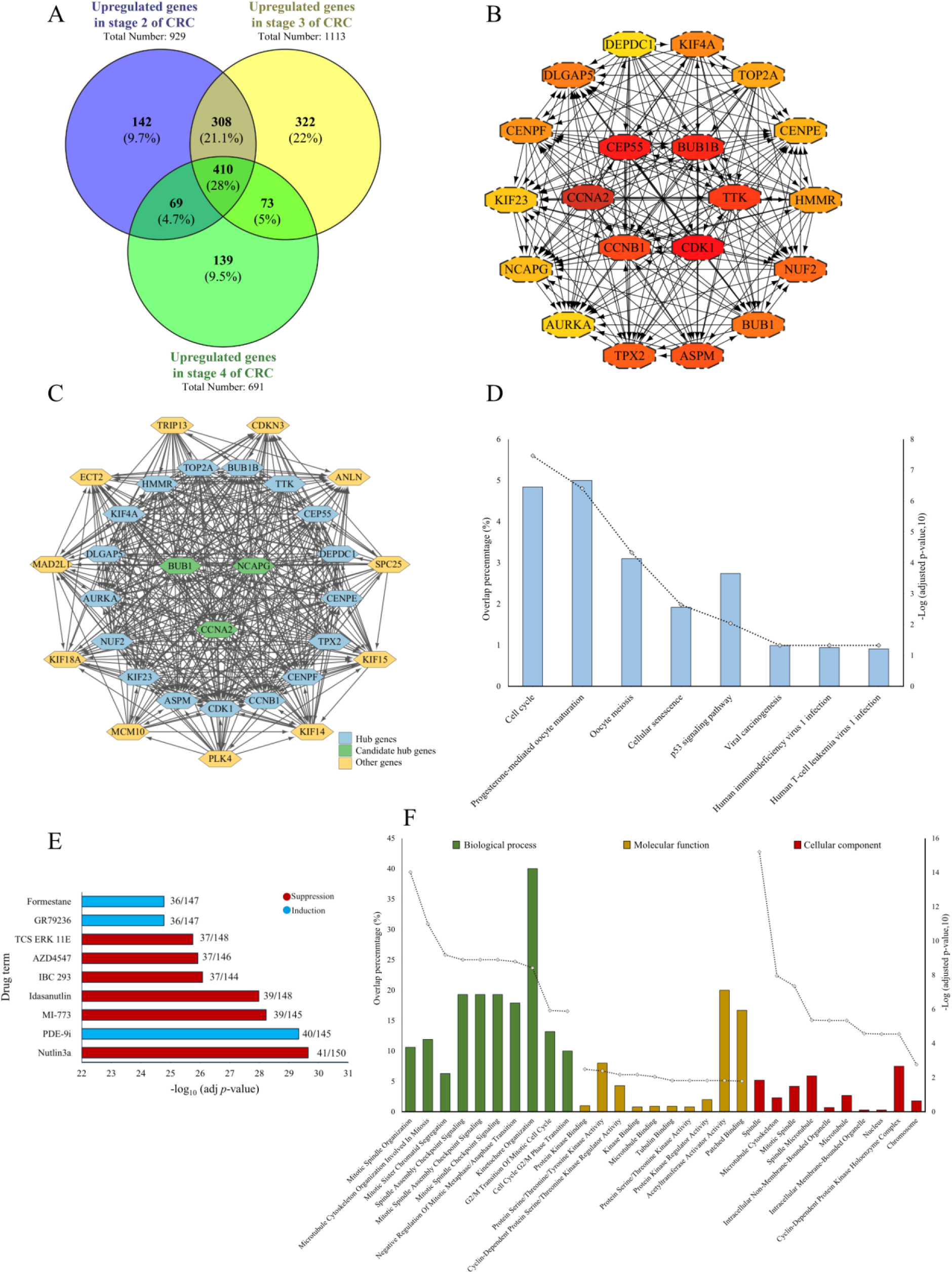
Integrated bioinformatic analysis of genes commonly upregulated across CRC stages. **(A)** Venn diagram of genes upregulated in stage II (929), stage III (1,113) and stage IV (691) CRC relative to normal tissue, percentages are of the 1,463 genes upregulated in at least one stage; **(B)** PPI sub-network of the 20 hub genes ranked highest by MCC among the 378 protein-coding genes upregulated at all three stages. Node colour reflects MCC rank, with darker red indicating higher rank; **(C)** Highest-scoring module extracted from the full PPI network with MCODE (31 nodes, 414 edges, score = 27.6). Blue, hub genes; green, candidate hub genes; yellow, additional module members; **(D)** KEGG pathway enrichment of the 410 commonly upregulated genes; **(E)** Drug-gene enrichment ranked by −log₁₀ (adjusted p); red, suppression; blue, induction. Numbers indicate overlapping genes per total targets in each drug set; **(F)** GO enrichment separated into biological process, molecular function and cellular component. In **(D)** and **(F)**, bars indicate the percentage of gene overlap (left axis), and the dotted line indicates −log₁₀ (adjusted p) (right axis).

Of the 410 shared genes, 378 were protein-coding and mapped onto a PPI network. The 20 highest- ranking nodes were designated hub genes using MCC algorithm **(Figure 1B)**. These included *CCNB1*, *CDK1*, *BUB1*, *CCNA2*, *NCAPG* and *CEP55*, which function in cell-cycle progression, mitotic spindle assembly and chromosome segregation. MCODE decomposition of the same network returned a single highest-scoring module (score = 27.6) comprising all 20 hub genes together with 12 additional nodes **(Figure 1C)**.

KEGG enrichment of the 410 shared genes returned cell cycle as the most significantly enriched pathway (adjusted p = 9.59E-15), followed by some other crucial pathways in cancer such as cellular senescence and p53 signalling **(Figure 1D)**. The oocyte-associated terms, as some other enriched terms, share their annotated members with the mitotic machinery and are therefore driven by the same gene set rather than by tissue-specific biology.

Drug-gene enrichment against the Proteomics Drug Atlas 2023 library prioritized nine compounds **(Figure 1E)**. Suppressive associations were dominated by MDM2 antagonists, nutlin-3a (41/150 genes), MI-773 (39/145) and idasanutlin (39/148), together with IBC-293, AZD4547 and TCS ERK 11E. That is while inductive associations comprised PDE-9i (40/145), formestane and GR79236 (36/147 each). GO analysis reinforced this mitotic signature **(Figure 1F)**. The most significantly enriched biological processes were mitotic spindle organization, microtubule cytoskeleton organization involved in mitosis and mitotic sister chromatid segregation; the leading molecular functions were protein kinase binding and patched binding; and the primary cellular components were spindle and mitotic microtubule cytoskeleton.

### 3.2 The hub gene set is enriched for targets of cell-cycle transcription factors such as NFY and the miR-192/miR-215 seed family

Gene set enrichment analysis (GSEA) against ENCODE ChIP-seq–derived target sets identified seven protein transcription factors significantly over-represented among the 20 hub genes (adjusted p < 0.05; **Table 3**). E2F4 ranked first (adjusted p = 1.00 × 10⁻²⁰), overlapping 17 hub genes including *TOP2A*, *BUB1B*, *NCAPG*, *CCNA2* and *CDK1*. FOXM1, a master regulator of the G2/M transition, overlapped 9 hub genes (adjusted p = 3.49 × 10⁻¹⁵), including *CCNA2*, *ASPM* and *KIF23*; of all regulators tested, FOXM1 showed the highest target-set specificity, with 9.5% of its annotated targets present among the hub genes. SIN3A, the scaffold subunit of the SIN3–HDAC corepressor complex, overlapped 15 hub genes (adjusted p = 3.64 × 10⁻¹⁴). The α and β subunits of nuclear transcription factor Y overlapped 13 and 17 hub genes, respectively (adjusted p = 1.68 × 10⁻⁷ and 3.02 × 10⁻⁹), consistent with the CCAAT boxes present in the promoters of many G2/M genes; *NCAPG* was absent from both subunit target sets. FOS and IRF3 each overlapped 5 hub genes at substantially weaker significance (adjusted p = 2.9 × 10⁻³ and 3.0 × 10⁻³).

**Table 3.**
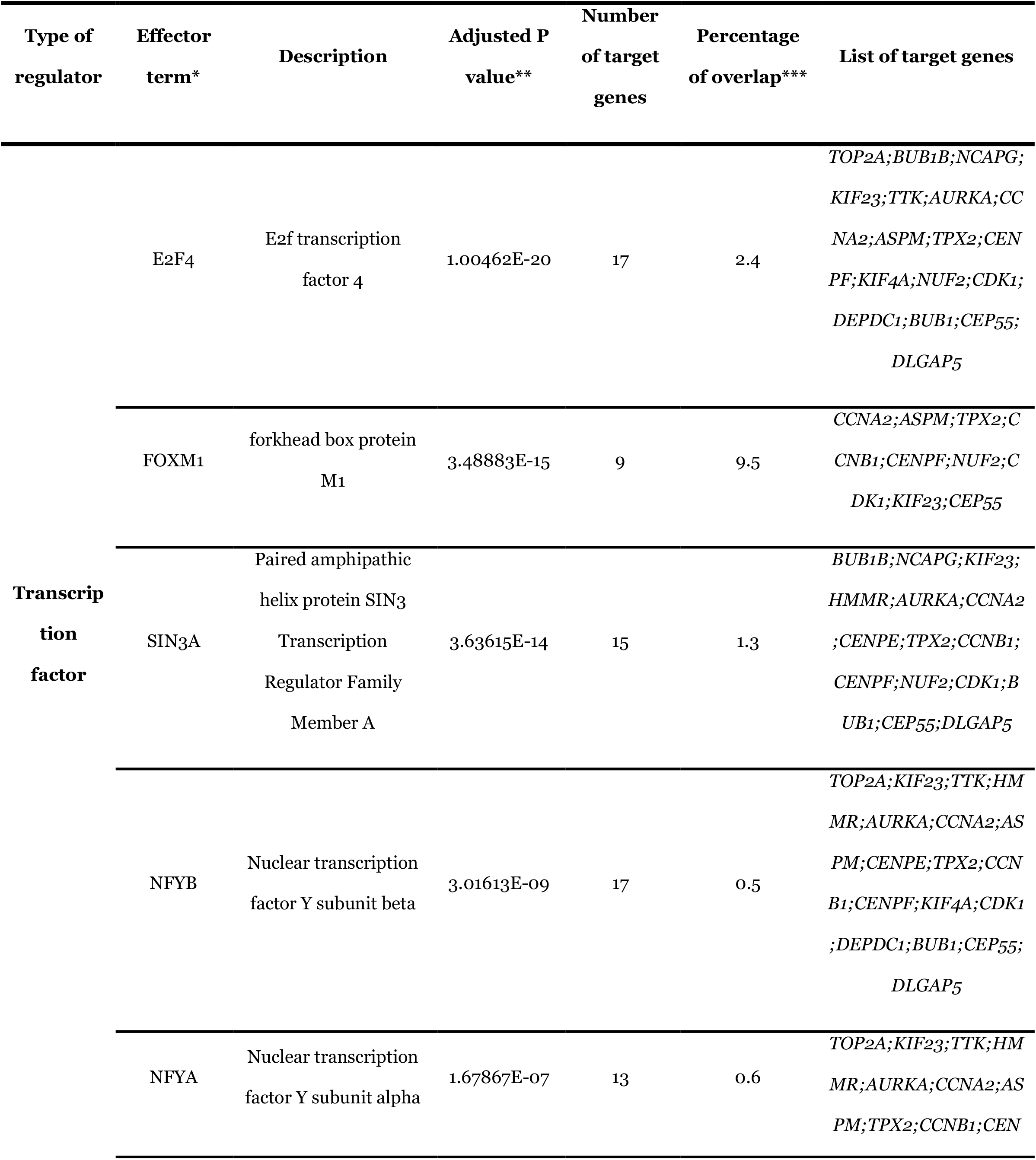
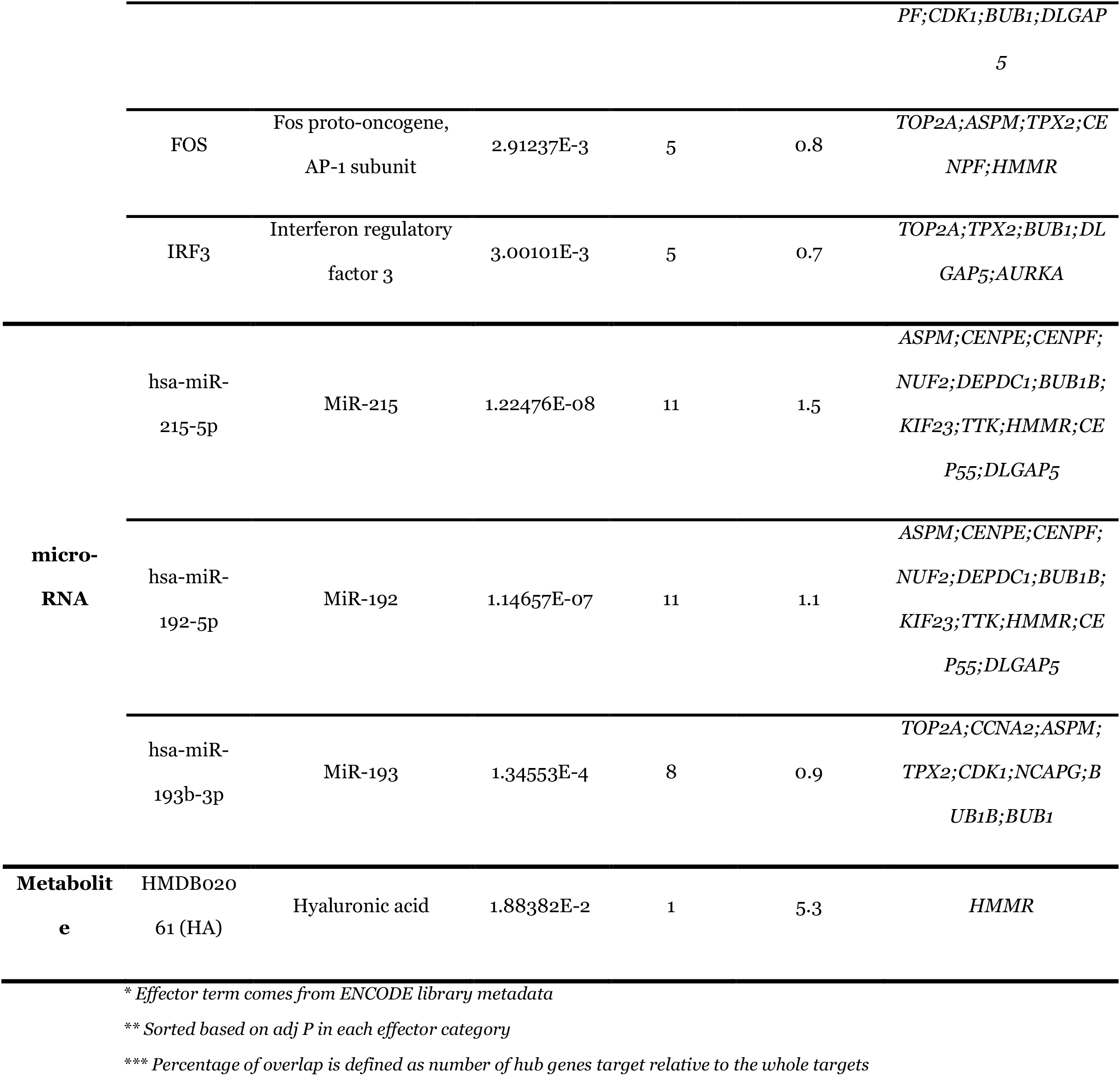
List of regulators associating with the expression level of the hub genes.

Among enriched miRNAs, hsa-miR-215-5p and hsa-miR-192-5p returned identical overlaps of 11 hub genes, including *ASPM*, *CENPF*, *NUF2* and *HMMR* (adjusted p = 1.22 × 10⁻⁸ and 1.15 × 10⁻⁷, respectively); these microRNAs share a seed sequence, and their results are therefore not independent observations. hsa-miR-193b-3p overlapped 8 hub genes, including *NCAPG* and *BUB1* (adjusted p = 1.35 × 10⁻⁴). Both miR-192 and miR-215 are reported to be downregulated in colorectal carcinoma [49], a direction compatible with derepression of the mitotic targets identified here.

A single metabolite, hyaluronic acid, reached significance (adjusted p = 1.9 × 10⁻²) through one hub gene, *HMMR*, which encodes the hyaluronan-mediated motility receptor. These analyses draw on reference ChIP-seq and miRNA–target databases rather than on regulator expression in the present cohort and therefore nominate candidate upstream regulators of the hub gene set rather than establishing regulatory relationships in CRC.

### 3.3 *In silico* functional characterization prioritizes *BUB1*, *CCNA2* and *NCAPG* among the hub genes

To assess whether the hub genes are functionally required in CRC, gene-level dependency data were retrieved from the DepMap 22Q4 genome-wide CRISPR–Cas9 screen for 59 colorectal cancer cell lines **(Figure 2A)**. Thirteen of the 20 hub genes reached a mean probability of dependency > 0.5 across the panel, indicating a common requirement for CRC cell proliferation or survival. *CDK1*, *TOP2A*, *BUB1*, *TTK* and *KIF23* showed uniformly high dependency and formed a single terminal cluster, whereas *CEP55*, *ASPM*, *CENPF*, *HMMR*, *DEPDC1*, *DLGAP5* and *KIF4A* showed low or heterogeneous dependency, consistent with functional redundancy or context-dependent requirement.

**Figure 2.**
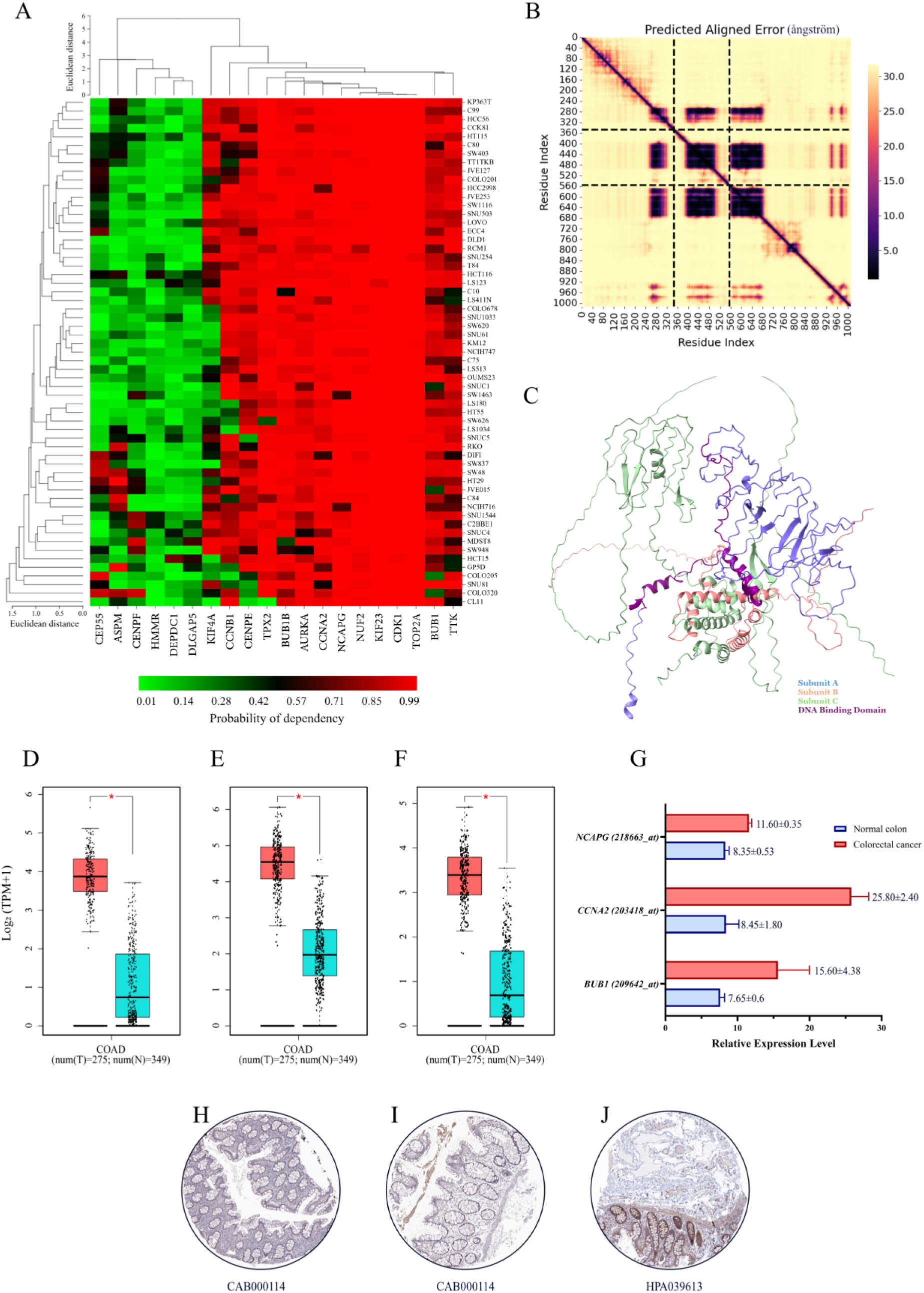
Functional validation of CRC hub genes using various bioinformatic methods. **(A)** Clustered heatmap showing gene dependency scores of the 20 identified hub genes across 59 CRC cell lines, derived from the DepMap 22Q4 CRISPR-Cas9 knockout screening dataset. Red indicates high dependency (greater essentiality for cell viability), while green denotes low dependency. Hierarchical clustering of both genes and cell lines was performed using Euclidean distance ranging from 0 to 6, and 0 to 1.5 for genes and cell lines, respectively; **(B)** Predicted Aligned Error plot for interaction of NFY subunits according to AF3; **(C)** Predicted structure and domain organization of NFY complex; **(D-F)** Box plots depicting expression levels of *BUB1*, *CCNA2*, and *NCAPG* in colorectal adenocarcinoma (COAD) versus normal colon tissue based on GEPIA analysis of TCGA and GTEx datasets (275 tumor vs. 349 normal samples). Adjusted *P* value < 0.05, indicated by a single red asterisk); **(G)** Bar plot from BioGPS illustrating relative expression levels of the same genes in healthy colon versus CRC samples; **(H-I)** antibody IDs in the figure; **(H)** IHC of CCNA2 in normal colon tissue; **(I)** IHC of CCNA2 in colon adenocarcinoma tissue; **(J)** IHC of NCAPG in normal colon tissue. Statistical significance: *P value < 0.05.

After taking all data into account and an accurate consideration of upstream regulatory factors while combining the results with literature mining, 3 (*BUB1*, *CCNA2* and *NCAPG*) of 20 upregulated hub genes were opted for further downstream experiments. Among the seven enriched transcription factors **(Table 3)**, NF-YA and NF-YB were the only regulators whose target sets contained both *BUB1* and *CCNA2* while excluding *NCAPG*; E2F4 and SIN3A target all three. Because NF-Y functions only as an obligate NF-YA–NF-YB–NF-YC heterotrimer, the architecture of the complex and its subunit interfaces were modelled with AlphaFold 3 (AF3) **(Figure 2B, C)**. The model reproduced the expected histone-fold dimer of NF-YB and NF-YC with inter-subunit placement; the N- and C-terminal segments of NF- YA and NF-YC were predicted with low confidence and high predicted aligned error, consistent with their annotated intrinsic disorder (ipTM = 0.41, pTM = 0.35).

All three genes were significantly overexpressed in colon adenocarcinoma relative to non-tumour colon tissue in GEPIA (TCGA + GTEx; 275 tumour, 349 normal; one-way ANOVA, q < 0.05; **Figure 2D–F**), with median log₂ (TPM + 1) values of 3.9 vs 0.9 for *BUB1*, 4.5 vs 2 for *CCNA2* and 3.5 vs 0.5 for *NCAPG*. The same direction of change was observed in the expression dataset of GeneAtlas U133A, gcrma **(Figure 2G)**, where relative expression changed significantly for *BUB1 (209642_at)*, *CCNA2 (203418_at)* and *NCAPG (218663_at)* about 2-, 3-and 1.4-fold change, respectively. Protein-level expression was examined in the Human Protein **(Figure 2H–J)**. CCNA2 showed strong nuclear staining in a minority (< 25%) of enterocytes in normal colon, with no detectable signal in endothelial, stromal or lymphoid cells **(Figure 2H)**. In colon adenocarcinoma, CCNA2 showed strong nuclear immunoreactivity in tumour epithelium **(Figure 2I)**. NCAPG showed moderate-to-weak cytoplasmic and membranous staining in > 75% of glandular cells **(Figure 2J)**, a distribution consistent with the cytoplasmic sequestration of condensin I during interphase. These results are single representative cores per condition and are not quantitative.

Finally, gene-disease associations were retrieved from DisGeNET v25.1 for the concept "colorectal carcinoma". Nineteen of the 20 hub genes had at least one curated association **(Table 4)**; *DEPDC1* had none. Disease Specificity Index (DSI) values, which increase as a gene is associated with fewer diseases, ranged from 0.475 (*AURKA*) to 0.743 (*NCAPG*), and Disease Pleiotropy Index (DPI) values from 0.269 (*NCAPG*) to 0.808 (*CDK1*, *BUB1B*). *CDK1*, *BUB1B* and *TOP2A* comprised the largest numbers of supporting publications with the lowest DSI and highest DPI of the set, identifying them as broadly pleiotropic rather than CRC-selective. *NCAPG* showed the opposite profile, the highest DSI and lowest DPI, indicating comparatively selective disease association.

**Table 4.** Gene-disease association for detected hub genes and CRC.

| **Gene symbol** | **Gene full name** | **Protein class** | **DSI** | **DPI** | **Number of PubMed articles supporting GDA** | **First and last year of GDA report** |
| --- | --- | --- | --- | --- | --- | --- |
| ***CDK1*** | Cyclin dependent kinase 1 | Kinase | 0.482 | 0.808 | 11 | 1995-2019 |
| ***CEP55*** | Centrosomal protein 55 | Protein subunit | 0.534 | 0.731 | 1 | 2006 |
| ***BUB1B*** | BUB1 mitotic checkpoint serine/threonine kinase | Kinase | 0.502 | 0.808 | 4 | 2002-2016 |
| ***CCNA2*** | Cyclin A2 | Enzyme modulator | 0.578 | 0.654 | 3 | 2012-2017 |
| ***TTK*** | TTK protein kinase | Kinase | 0.555 | 0.731 | 3 | 2011-2019 |
| ***CCNB1*** | Cyclin B1 | Enzyme modulator | 0.512 | 0.731 | 8 | 1997-2017 |
| ***ASPM*** | Assembly factor for spindle microtubules | Protein subunit | 0.526 | 0.769 | 2 | 2013-2014 |
| ***TPX2*** | TPX2 microtubule nucleation factor | Cellular structure protein | 0.541 | 0.692 | 3 | 1999-2015 |
| ***NUF2*** | NUF2 component of NDC80 kinetochore complex | Cellular structure protein | 0.678 | 0.346 | 1 | 2014 |
| ***BUB1*** | BUB1 mitotic checkpoint serine/threonine kinase | Kinase | 0.528 | 0.731 | 10 | 2000-2018 |
| ***DLGAP5*** | DLG associated protein 5 | Receptor | 0.678 | 0.423 | 2 | 2012-2019 |
| ***KIF4A*** | Kinesin family member 4A | Cellular structure protein | 0.647 | 0.462 | 2 | 2018 |
| **CENPF** | Centromere protein F | Protein subunit | 0.601 | 0.654 | 1 | 2018 |
| **HMMR** | Hyaluronan mediated motility receptor | Receptor | 0.544 | 0.615 | 2 | 2008, 2017 |
| **TOP2A** | DNA topoisomerase II alpha | Enzyme | 0.497 | 0.731 | 5 | 1999-2007 |
| **CENPE** | Centromere protein E | Cellular structure protein | 0.636 | 0.692 | 1 | 2013 |
| **NCAPG** | Non-SMC Condensin I Complex Subunit G | Complex subunit | 0.743 | 0.269 | 2 | 2018, 2022 |
| **KIF23** | Kinesin family member 23 | Cellular structure protein | 0.682 | 0.385 | 1 | 2013 |
| **AURKA** | Aurora kinase A | Kinase | 0.475 | 0.731 | 15 | 2004-2019 |
| **DEPDC1** | DEP domain containing 1 | Nucleic acid binding | 0.670 | 0.577 | 0 | n.a. |

### 3.4 Melatonin reduces CRC cell viability and induces apoptosis

Sw480 cells were exposed to melatonin (0–20 mM) for 48 h and viability assessed by MTT assay **(Figure 3A)**. Viability was relatively unchanged between 0.63 and 1.25 mM, declined to 55.1% at 2.50 mM, and fell below 3% at concentrations of 5 mM and above. Four-parameter logistic regression of the log-transformed data returned an IC₅₀ of 2.63 mM (95% CI= 0.81–8.84; Hill slope = -5.22; R² = 0.99; **Figure 3B)**. Two different concentrations spanning the transition region of the curve, IC₃₅ (2.33mM) and IC₆₅ (2.91 mM), were used for all subsequent experiments.

**Figure 3.**
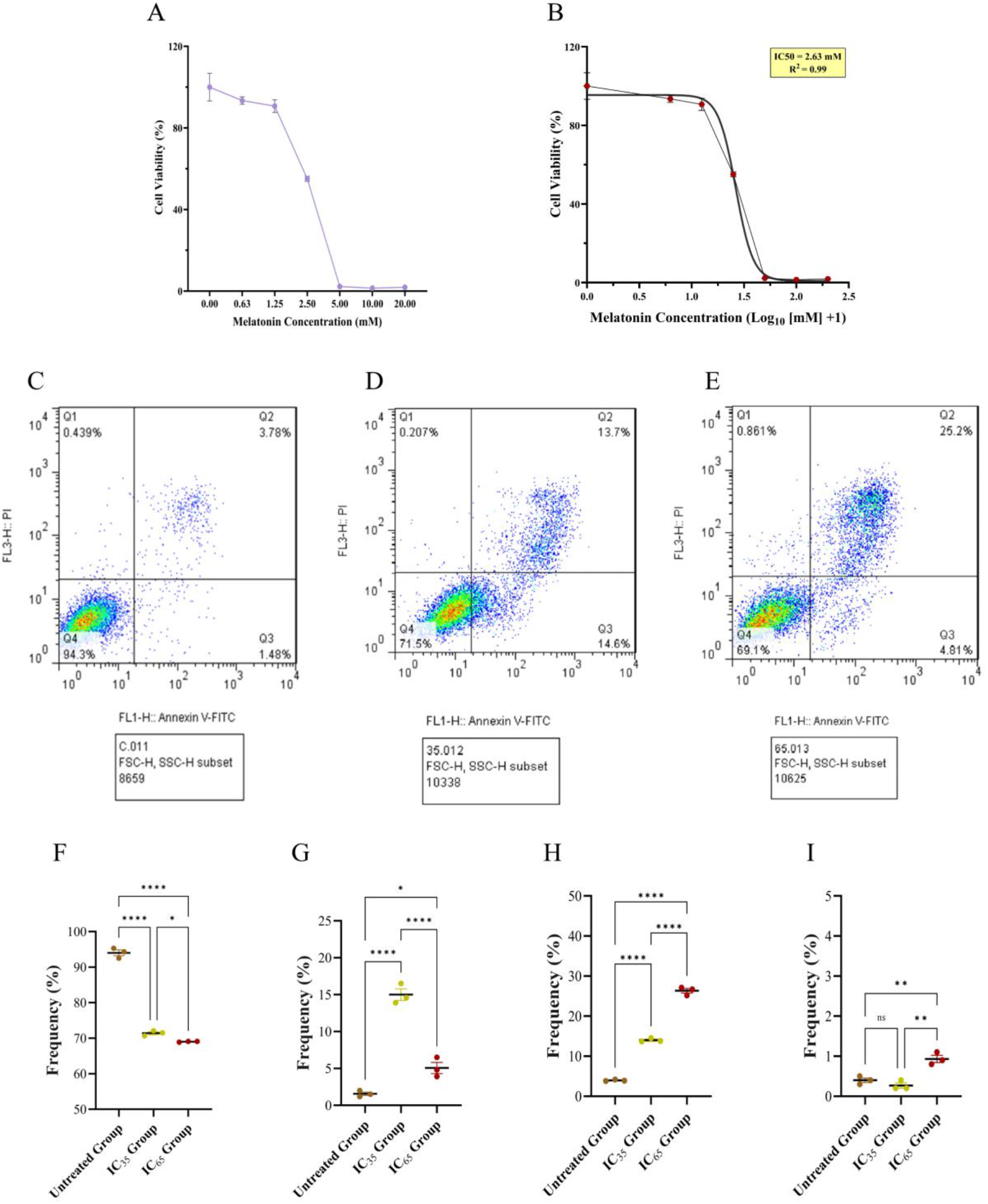
Melatonin reduces viability and induces apoptosis in SW480 cells. **(A)** Viability after 48 h of melatonin exposure (0–20 mM), measured by MTT assay. Mean ± SD; n = 5 independent experiments. Concentrations are plotted as evenly spaced categories; **(B)** Four-parameter logistic fit to the log₁₀-transformed data. IC₅₀ = 2.63 mM (95% CI 0.81–8.84); Hill slope = -5.22; R² = 0.99. Top and bottom were constrained. **(C– E)** Representative annexin V–FITC/PI dot plots for untreated **(C)**, IC₃₅ **(D)** and IC₆₅ **(E)** cultures after 48 h; Events analysed in one replicate as a sample: 8,659 **(C)**, 10,338 **(D)**, 10,625 **(E)**, after gating on FSC-H/SSC-H; **(F–I)** Quantification of **(F)** viable (Q4), **(G)** early apoptotic (Q3), **(H)** late apoptotic (Q2) and **(I)** PI-only (Q1) fractions. Individual values with mean ± SD; n = 3 independent experiments. One-way ANOVA with post-hoc Tukey: ns, not significant; *P < 0.05; **P < 0.01; ****P < 0.0001.

Apoptosis was quantified by annexin V–FITC/PI staining after 48 h **(Figure 3C–E)**, with populations resolved as viable (annexin V⁻/PI⁻, Q4), early apoptotic (annexin V⁺/PI⁻, Q3), late apoptotic (annexin V⁺/PI⁺, Q2) and necrotic (annexin V⁻/PI⁺, Q1). Untreated cultures were 94.03±1.42% viable with 1.56±0.40% only-annexin V⁺ events (n =3; **Figure 3F–I**). Melatonin reduced the viable fraction to 71.43±0.70% at IC₃₅ and 68.23±0.11% at IC₆₅ (both P < 0.0001 versus untreated; One-way ANOVA with post-hoc Tukey), with a corresponding rise in only-annexin V⁺ events to 14.04±0.43% and 5.06±1.32%.

The two concentrations differed in the distribution of apoptotic cells rather than in the total. At IC₃₅, early and late apoptotic fractions were comparable (15.01±1.34% and 14.04±0.43), whereas at IC₆₅ the late apoptotic fraction increased to 25.83±1.01% (P < 0.0001) and the early apoptotic fraction fell to 5.06±1.32% (P < 0.0001), consistent with progression of a similarly sized apoptotic population to a later stage. PI-only events remained below 1% in all conditions but increased at IC₆₅ relative to both untreated and IC₃₅ cultures (0.41±0.10% versus 0.26±0.11% and 0.93±0.15%; P < 0.01)

### 3.5 Melatonin represses the NF-Y target genes *BUB1* and *CCNA2* but not *NCAPG*

To test whether the NF-Y–target relationships predicted in section 3.2 are sensitive to melatonin, transcript levels of *BUB1*, *CCNA2* and *NCAPG* were measured by qPCR after 48 h of treatment at the IC₃₅ and IC₆₅ concentrations (n = 3 independent experiments each having 3 technical replicates; **Figure 4A–C, E–G**).

**Figure 4.**
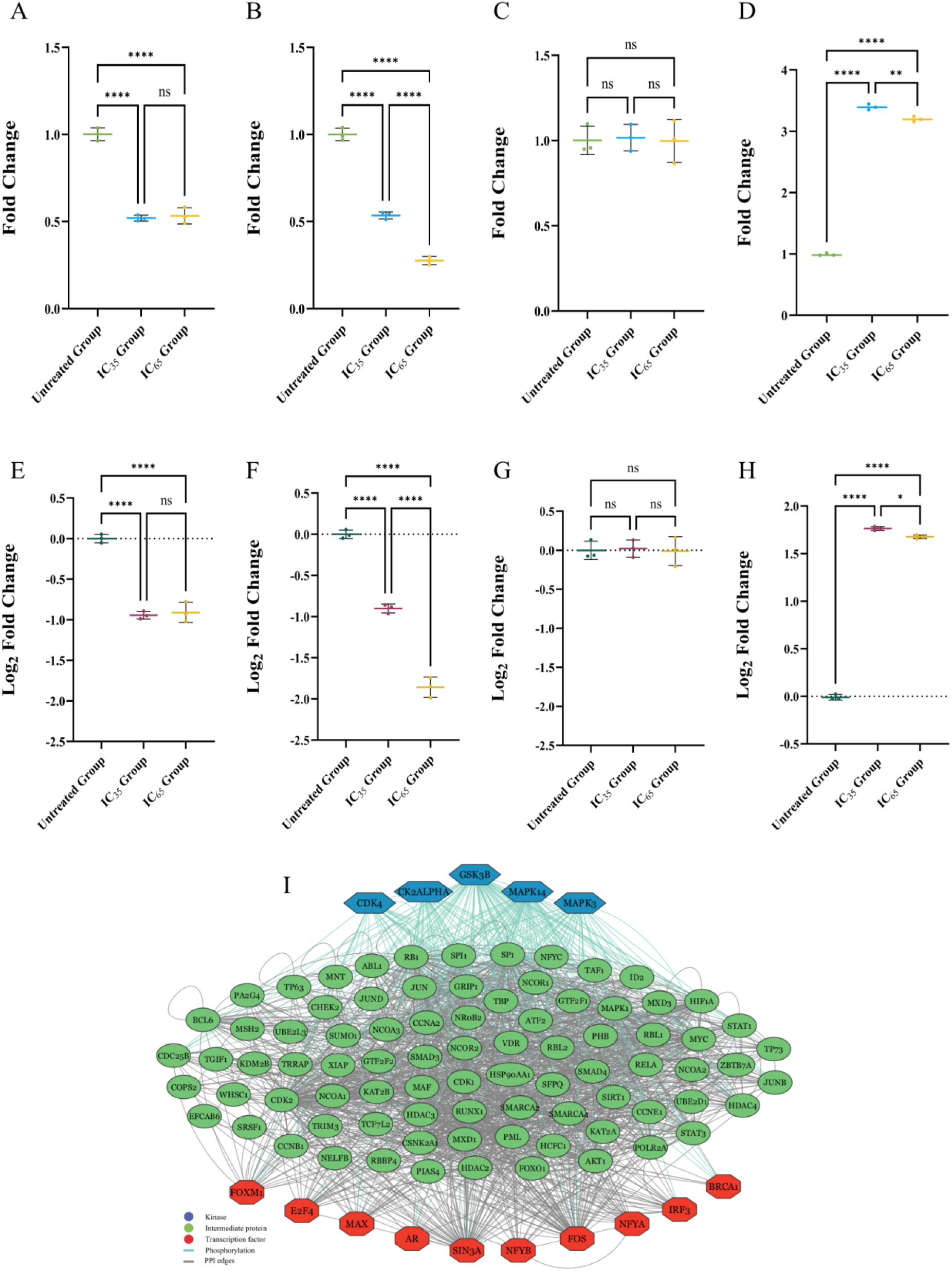
Melatonin represses *BUB1* and *CCNA2* and spares *NCAPG* while inducing *B2M*. **(A–D)** Relative transcript levels (fold change) and **(E–H)** the same data as log₂ fold change for **(A, E)** BUB1, **(B, F)** CCNA2, **(C, G)** NCAPG and **(D, H)** B2M, after 48 h of melatonin treatment at the IC₃₅ and IC₆₅ concentrations. Values were normalized to the geometric mean of *GAPDH* and *ACTB* by the Pfaffl method and expressed relative to untreated cultures. Individual values with mean ± SD; n = 3 independent biological replicates. One-way ANOVA with post hoc Tukey applied to fold change values: ns, not significant; *P < 0.05; **P < 0.01; ****P < 0.0001. **(I)** Upstream regulatory network for the 20 hub genes reconstructed with X2K. Blue hexagons: kinases; green ellipses: intermediate proteins (PPI expansion); red hexagons: transcription factors (ChEA/ENCODE). Teal edges, phosphorylation; grey edges, protein–protein interactions. The hub genes themselves are not represented as nodes. Self-loops are either autophosphorylation or self-interaction.

*BUB1* transcript fell to 0.52 ± 0.02 of the untreated level at IC₃₅ and 0.53 ± 0.05 at IC₆₅ (both *P* < 0.0001), with no difference between the two concentrations (*P* = 0.89). *CCNA2* fell to 0.54 ± 0.02 at IC₃₅ and further to 0.28 ± 0.02 at IC₆₅ (all pairwise comparisons *P* < 0.0001), indicating a concentration-dependent response over this range. *NCAPG*, which was absent from both the NF-YA and NF-YB target sets according to bioinformatic data mining **(Table 3)**, was unchanged at either concentration (*P* = 0.98 and 0.99).

Among the seven transcription factors enriched in the hub gene set, NF-YA and NF-YB were the only ones whose target sets include *BUB1* and *CCNA2* while excluding *NCAPG*; E2F4 and SIN3A target all three, and FOXM1, FOS and IRF3 target at most one **(Table 3)**. The observed expression pattern is therefore compatible with reduced NF-Y-dependent transcription, although three genes cannot discriminate between this and other regulators sharing the same target configuration.

Three candidate reference genes were evaluated. *B2M* transcript increased 3.19 ± 0.05-fold at IC₃₅ and 3.15 ± 0.02-fold at IC₆₅ relative to untreated cultures (both *P* < 0.0001; **Figure 4D, H**) and was therefore excluded as unstable under treatment. *GAPDH* and *ACTB* Ct values varied by less than 0.5 cycles across conditions (M-value < 1.5), and their geometric mean was utilized for normalization.

Independently, the 20 hub genes were submitted to X2K to reconstruct an upstream regulatory network **(Figure 4I)**. The transcription factor layer comprised 98 nodes, including NFYA and NFYB alongside E2F4, FOXM1, SIN3A, FOS and IRF3, the same factors recovered in section 3.2, since X2K queries overlapping ChIP-seq libraries. NFYA and NFYB ranked 5 and 8 by degree among 10 transcription factor nodes. The protein kinase layer comprised CDK4, CSNK2A1, GSK3B, MAPK3 and MAPK14, connected to the transcription factor layer through 83 intermediate proteins. This network is derived from reference interaction databases and is independent of the melatonin treatment.

### 3.6 Melatonin arrests CRC cells in G1 and impairs cell proliferation and migration

Cell-cycle distribution was determined by propidium iodide DNA content analysis after 48 h of melatonin treatment and modelled with the Watson pragmatic algorithm (FlowJo v10; **Figure 5A–F**). The G2:G1 mean fluorescence ratio was 1.94–1.95 across all samples and G1 coefficients of variation were 4.99–6.03, confirming acceptable staining and resolution.

**Figure 5.**
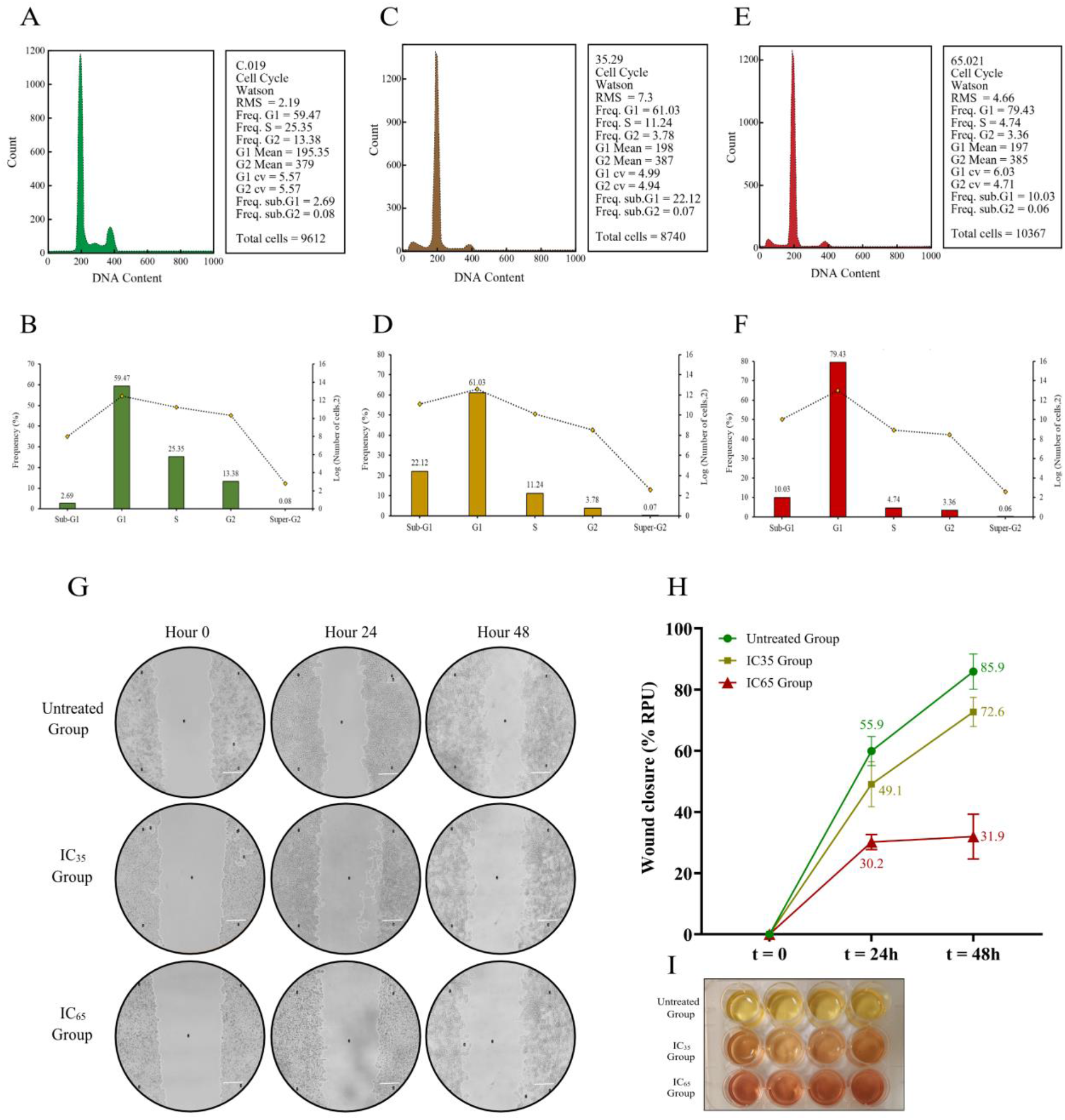
Melatonin arrests CRC cells in G1 and slows wound closure. **(A, C, E)** DNA content histograms for untreated **(A)**, IC₃₅-treated **(C)** and IC₆₅-treated **(E)** cells after 48 h, with Watson pragmatic model fits. Events analysed in the representative sample: 9,612 **(A)**, 8740 **(C)**, 10,367 **(E)**. RMS goodness-of-fit: 2.19, 7.30, 4.66. **(B, D, F)** Corresponding distributions across sub-G1, G1, S, G2/M and >4N regions (bars, left axis; percentage of total events) with event counts per region (dotted line, right axis; log₂). n=3 independent replicates for this experiment; **(G)** Representative phase-contrast images of the scratch area at 0, 24 and 48 h. Scale bar, 150 µm; **(H)** Wound closure quantified in ImageJ v1.52 and expressed as relative pixel unit (RPU); mean ± SD, n = 3 independent experiments. Values above each series indicate the mean closure rate over 48 h; **(I)** Culture plates photographed at 48 h; yellow medium indicates acidification.

Untreated cultures showed an asynchronous profile with 59.47% of events in G1, 25.35% in S and 13.38% in G2/M, and 2.69% in the sub-G1 region **(Figure 5A, B)**. Melatonin reduced the S and G2/M fractions at both concentrations: to 11.24% and 3.78% at IC₃₅ **(Figure 5C, D)** and to 4.74% and 3.36% at IC₆₅ **(Figure 5E, F)**. Expressed as a proportion of the cycling (non-sub-G1) population, the G1 fraction rose from 60.6% in untreated cultures to 80.3% at IC₃₅ and 90.8% at IC₆₅, with corresponding decreases in S (25.8% → 14.8% → 5.4%) and G2/M (13.6% → 5.0% → 3.8%), indicating concentration-dependent accumulation at G1 and depletion of cells beyond the G1/S boundary. The sub-G1 fraction increased to 22.12% at IC₃₅ and 10.03% at IC₆₅. This distribution is consistent with the transcriptional changes in section 3.5: cyclin A2, encoded by the melatonin-repressed *CCNA2*, is required for S-phase entry and progression, so its downregulation provides a candidate mechanism for the observed G1/S block.

Collective migration and proliferation were assessed by scratch assay over 48 h **(Figure 5G, H)**. Wound closure in untreated cultures reached 85.9% of the initial wound area by 48 h, compared with 72.6% at IC₃₅ and 31.9% at IC₆₅ (n = 3 biological replicates each having 3 technical replicates). Closure at IC₆₅ was largely inhibited by even 24 h treatment while changing little thereafter (30.2 to 31.9 units), whereas untreated and IC₃₅ cultures continued to close between 24 and 48 h. Because melatonin reduced viability and arrested proliferation under these conditions, the assay reports the combined contribution of migration, division and survival, and does not isolate migratory capacity.

Consistent with reduced cell number and metabolic output, phenol red in the culture medium remained red in treated wells after 48 h while acidifying to yellow in untreated wells (qualitative observation; **Figure 5I**).

### 3.7 Melatonin treatment increases cellular fluorescence and produces a non-monotonic change in DCF signal

SW480 cells were treated for 48 h and imaged by Epifluorescence (Ex 405 nm/Em 455) to assess intracellular fluorescence change upon melatonin treatment **(Figure 6A)**. CTCF increased from 56.72 ± 11.23 RFU in untreated cultures to 62.44 ± 9.51 RFU at IC₃₅ and 71.51 ± 14.47 RFU at IC₆₅ (mean ± SD; n = 270 cells from 3 independent experiments; One-way ANOVA post-hoc Tukey; **Figure 6B**). Bright-field images showed cell rounding and reduced density in both treated groups, consistent with the viability and cell-cycle data in sections 3.4 and 3.6.

**Figure 6.**
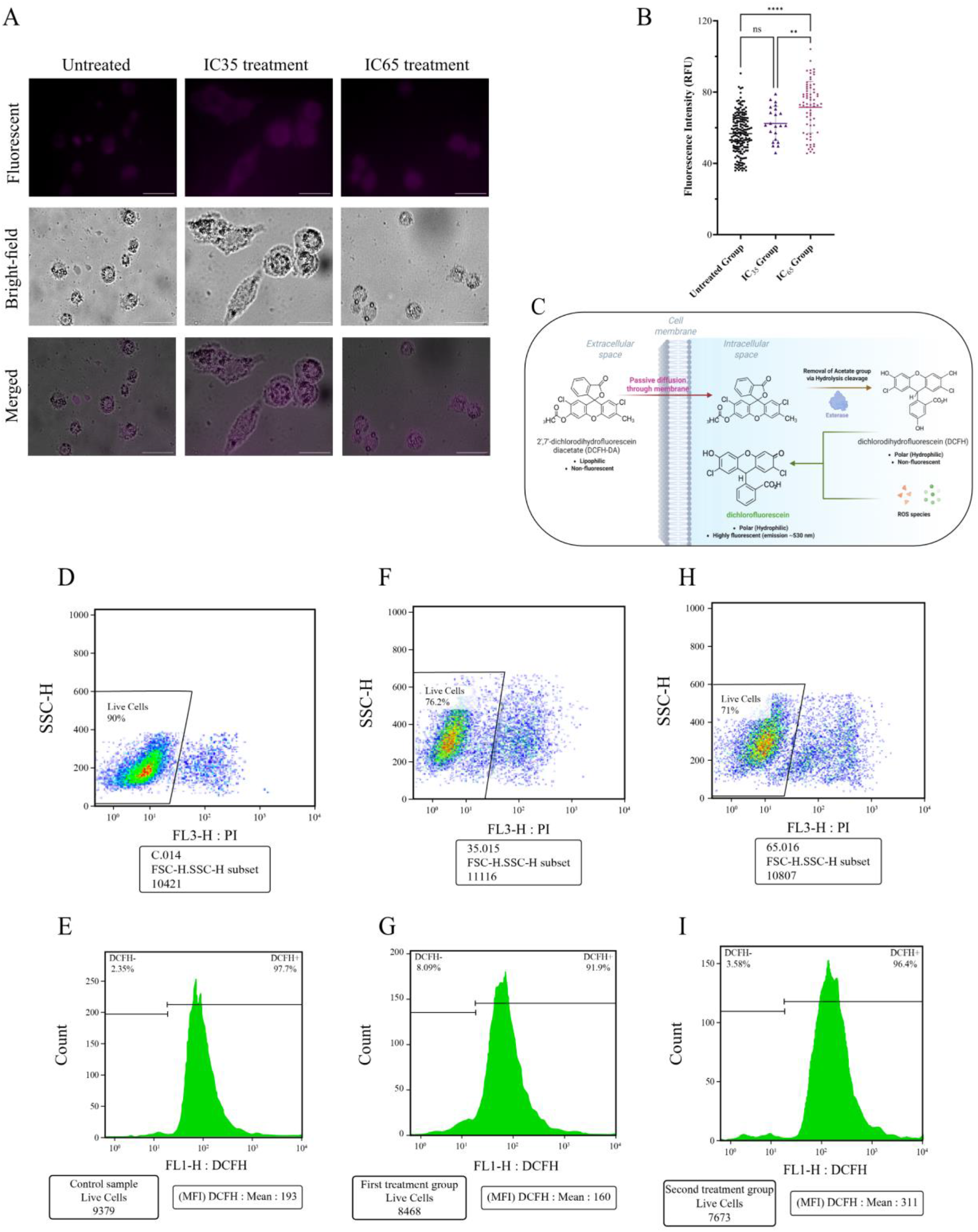
Cellular fluorescence and DCF signal after melatonin treatment. **(A)** Representative fluorescence (Ex 405 nm / Em 455 nm), bright-field and merged images after 48 h of treatment. Objective, 40×; identical acquisition settings for all conditions. Scale bar, 5 µm. **(B)** CTCF quantification (relative fluorescence units). Each point represents one cell; n = 180 (untreated), 22 (IC₃₅) and 68 (IC6₅) cells from 3 independent experiments. Mean ± SD; One-way ANOVA with post-hoc Tukey: ns, not significant; **P < 0.01; ****P < 0.0001. (C) Principle of DCFH-DA detection. Created with BioRender.com (https://app.biorender.com/illustrations/68173a491aaff24f1b764221). **(D, F, H)** PI versus SSC-H plots defining the live-cell gate for untreated **(D)**, IC₃₅ **(F)** and IC₆₅ **(H)** cultures; live/total events 9379/10421, 8468/11116 and 7673/10807; **(E, G, I)** DCF fluorescence histograms for the gated live cells in each condition, with mean fluorescence intensity of 193, 160 and 311. Gate boundaries and instrument settings were identical across samples.

Intracellular oxidant levels were measured with 2′,7′-dichlorodihydrofluorescein diacetate (DCFH-DA), which is hydrolysed by cellular esterases to non-fluorescent DCFH and oxidized to fluorescent DCF **(Figure 6C)**. Analysis was restricted to the PI-negative population, which comprised 90.0% of events in untreated cultures (9,379 of 10,421), 76.2% at IC₃₅ (8,468 of 11,116) and 71.0% at IC₆₅ (7,673 of 10,807; **Figure 6D, F, H**); values consistent with the annexin V data in section 3.4.

Within this population, mean DCF fluorescence was 193 in untreated cultures, decreased to 160 at IC₃₅ and increased to 311 at IC₆₅ **(Figure 6E, G, I)**. Expressed as mean fluorescence intensity, the corresponding values were 193, 160 and 311, i.e. 0.83-fold and 1.61-fold relative to untreated cultures. This non-monotonic pattern might be because melatonin may act as a radical scavenger at lower concentrations and promotes oxidant accumulation in tumour cells at higher ones, although the present data are from a single acquisition per condition and DCF oxidation is not specific to any individual reactive species.

### 3.8 A proposed model for melatonin-mediated suppression of NF-Y target genes

Three observations from the preceding sections converge on nuclear transcription factor Y as a candidate mediator. First, NF-YA and NF-YB were among the seven transcription factors enriched in the target-set analysis of the 20 hub genes and were the only enriched factors whose target sets contain both *BUB1* and *CCNA2* while excluding NCAPG **(Table 3)**. Second, melatonin repressed *BUB1* and *CCNA2* while leaving *NCAPG* unchanged **(Figure 4A–C, E–G)**, the pattern predicted by that target configuration. Third, the resulting G1 accumulation and depletion of S-phase cells **(Figure 5)** is consistent with loss of cyclin A2, which is required for S-phase progression.

On this basis we propose that melatonin reduces NF-Y-dependent transcription of a subset of mitotic genes in CRC cells **(Figure 7)**. Melatonin is a small lipophilic molecule that enters cells by passive diffusion, and the concentrations used here (millimolar) exceed the affinity of MT1 and MT2 by several orders of magnitude, so a receptor-independent mechanism is the more plausible of the two possibilities; however, receptor involvement was not tested. The proposed model remains open for evaluation as some other experimental validation like changes in protein levels, ChIP–qPCR, loss- or gain-of-function experiments would be required for final verification of bioinformatic and preliminary findings.

**Figure 7.**
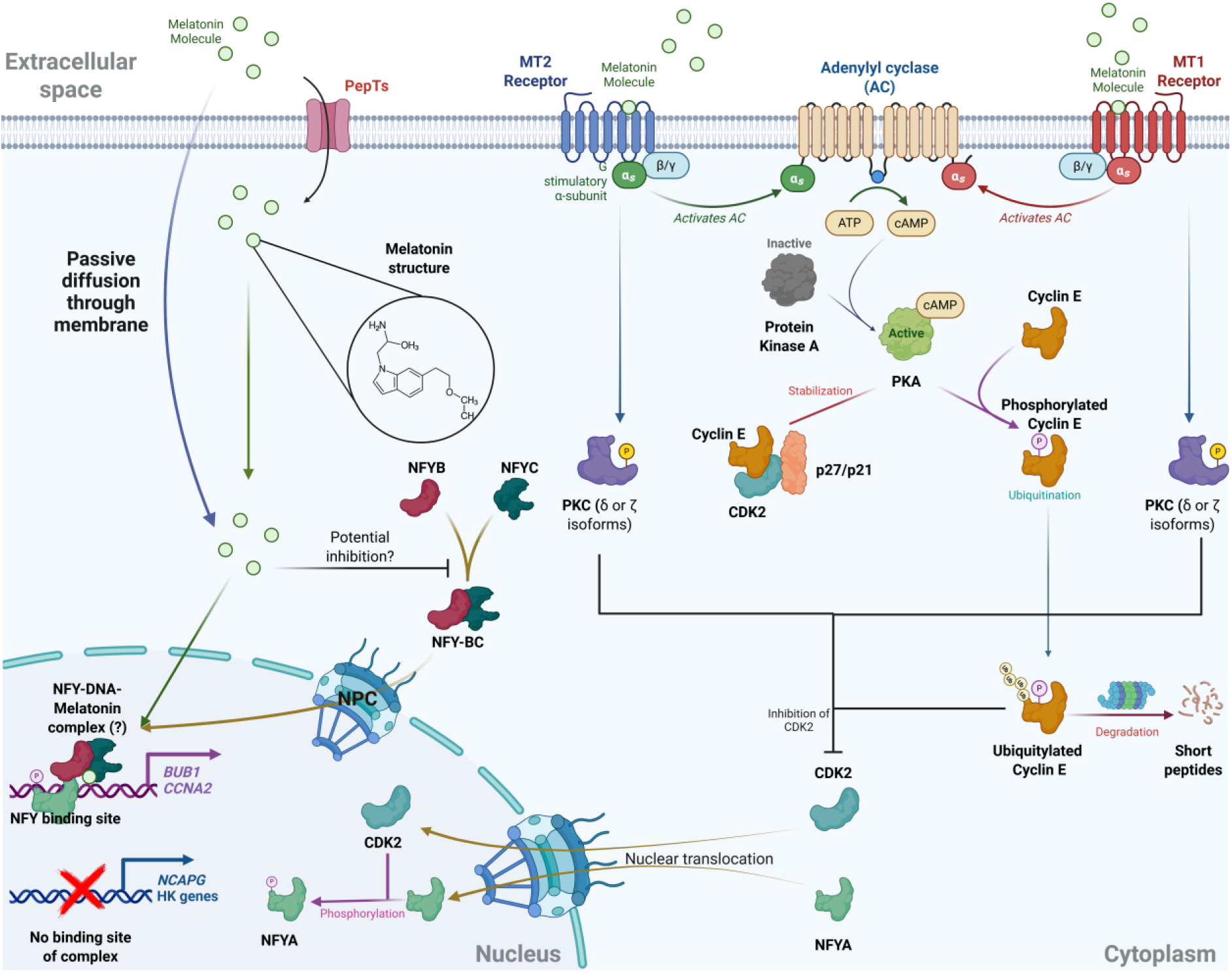
Proposed model for melatonin-mediated suppression of NF-Y target genes in CRC cells. NF-YB and NF-YC heterodimerize through their histone-fold domains in the cytoplasm and translocate to the nucleus, where NF-YA joins to form the DNA-binding trimer at CCAAT boxes. We hypothesize that melatonin reduces NF-Y-dependent transcription of *BUB1* and *CCNA2*, but not of *NCAPG*, which lacks an annotated NF-Y site. Created with BioRender.com (https://app.biorender.com/illustrations/681da9adf7d21bc61c1d9a23)

## 4. Discussion

The gene expression analysis in this study revealed that stage III of CRC exhibits the highest number of differentially expressed genes compared to normal tissue, while stage IV shows the lowest, and stage II standing intermediate. This pattern suggests a peak in transcriptional dysregulation during stage III, potentially reflecting heightened tumour progression and immune responses. Previous studies have similarly observed increased DEGs in stage III CRC, indicating significant biological alterations at this stage [50]. Conversely, the reduced number of DEGs in stage IV may result from tumour dedifferentiation and adaptation to metastatic environments, leading to a more uniform gene expression profile [51]. Moreover, our analysis revealed that approximately one-third of all upregulated genes (410 in total) across different stages of colorectal cancer are consistently overexpressed, suggesting a core set of genes pivotal to CRC pathogenesis. This observation aligns with prior studies identifying common upregulated genes across CRC stages, such as the 128 genes reported by Guiling Shi *et al.* [52].

The GSEA identified several key transcription factors, microRNAs, and a metabolite associated with the regulation of hub genes in CRC consistent with previous studies. Among all upstream TFs, E2F4 ranked first, overlapping 17 hub genes. E2F4 is the DNA-binding subunit of DREAM, which silences G2/M genes in quiescent cells, mutations of which have been described in CRC [53]. FOXM1, which displaces DREAM at those same promoters and promotes CRC growth and metastasis [54], showed the highest target-set specificity of any factor tested (9.5%). SIN3A, a distinct HDAC corepressor scaffold implicated in CRC progression [55], overlapped 15 hub genes. Regarding miRNAs, hsa-miR-215-5p has been reported to suppress tumour growth and metastasis in CRC by targeting genes involved in cell proliferation [56]. Similarly, hsa-miR-192-5p functions as a tumour suppressor in CRC, inhibiting cell proliferation and inducing apoptosis. The metabolite hyaluronic acid interacts with HMMR, influencing cell motility and proliferation. Elevated levels of HA and HMMR expression have been associated with CRC progression and poor prognosis [57].

Our results noted that Nutlin3a and PDE-9i are the most influential drugs on the pool of identified upregulated genes. Nutlin-3a restores p53 activity in CRC, inducing p21 and engaging the p53–p21– DREAM axis to repress G2/M drivers, potentially *CCNA2*, *BUB1*, and related mitotic module; thereby, enforcing arrest in HCT116 and allied models [58]. Beyond arrest, Nutlin-3a triggers ER-stress/CHOP-dependent DR5 upregulation and caspase-8 activation, sensitizing CRC cells to 5-FU and TRAIL independently of p53 status [59]. By contrast, PDE-9 inhibition elevates cGMP to activate PKG from a PDE9A-regulated pool, establishing a mechanistic route distinct from MDM2–p53 while remaining pharmacologically tractable [60].

The cytotoxicity of melatonin on the CRC cell line, as evidenced by an IC₅₀ of approximately 2.63 mM, also aligns with findings from other studies on melatonin cytotoxicity on other CRC cell lines. For example, LoVo colon cancer cells are reported to have an IC₅₀ of 2.1 mM for melatonin [61]. Moreover, in 5-FU-resistant SW480 cells, melatonin at concentrations of 1–2 mM significantly reduced thymidylate synthase (TYMS) expression, suggesting its potential to overcome chemoresistance [62]. However, these effective concentrations are substantially higher than physiological melatonin levels. In humans, nocturnal plasma melatonin concentrations typically range from 80 to 120 pg/mL (approximately 0.35 to 0.52 nM), while daytime levels drop to 10–20 pg/mL (around 0.04 to 0.09 nM) [63, 64]. This disparity underscores the challenge of achieving therapeutic melatonin concentrations *in vivo* without targeted delivery strategies. Nanotechnology-based approaches, such as selenium nanoparticles (SeNPs), have demonstrated improved anticancer effects and bioavailability of melatonin [65]. Additionally, pH-responsive sericin-based nanocarriers have been developed to co-deliver melatonin and resveratrol, showing enhanced cytotoxicity against breast cancer cells [66].

Our DEG analysis implies that melatonin selectively modulates specific transcriptional pathways, implicating the nuclear transcription factor Y complex as a central mediator as it is the only upstream TF indicated to influence *BUB1* and *CCNA2* expression, but not *NCAPG*. The NFY complex, comprising NFYA, NFYB, and NFYC subunits, binds to the CCAAT motif within promoter regions of various genes, including those pivotal for cell cycle progression such as *BUB1* and *CCNA2* [67]. NFY’s DNA-binding activity is contingent upon the phosphorylation status of NFYA, a process regulated by cyclin-dependent kinase 2 (CDK2) [68]. Melatonin’s capacity to inhibit CDK2 activity might potentially impede NFYA phosphorylation, hindering its nuclear translocation and subsequent DNA-binding ability.

On the other hand, melatonin can enter the cytoplasm via passive diffusion or through peptide transporters (PepTs) [69], as well as by binding to its high-affinity receptors, MT1 and MT2. Receptor-mediated activation leads to downstream signalling involving adenylyl cyclase, cAMP, and PKA, ultimately modulating Cyclin E–CDK2 complex formation and activity [13]. One critical consequence is the inhibition of CDK2, which may reduce NFYA phosphorylation, impairing its nuclear translocation and subsequent DNA-binding capability as part of the NFY heterotrimer (NFYA–NFYB–NFYC) [13]. Alternatively, melatonin may inhibit the nuclear import of the NFYB–NFYC dimer or disrupt NFY complex binding to the CCAAT promoter elements of NFY target genes. This would explain the selective repression of *CCNA2* and *BUB1*, whose promoters harbour functional NFY-binding sites, while *NCAPG*, lacking such a regulatory element, remains unaffected.

In this study, *β2M* exhibited significant upregulation upon melatonin treatment. Interestingly, the upregulation of *β2M* in response to melatonin treatment may have biological significance beyond its role as a housekeeping gene. *β2M* is a critical component of the major histocompatibility complex (MHC) class I molecules, which are essential for presenting endogenous antigens, including tumour-associated antigens and neoantigens, to cytotoxic CD8+ T cells [70]. Enhanced expression of *β2M* could potentially improve antigen presentation, thereby augmenting immune recognition and elimination of cancer cells.

Melatonin’s capacity to traverse cellular membranes via passive diffusion has been well-documented, primarily due to its amphiphilic nature and low molecular weight, which enable it to readily permeate lipid bilayers without requiring transporters or receptors [71]. This passive diffusion mechanism allows melatonin to accumulate intracellularly, acting directly within cytosolic and organelle compartments, a property that distinguishes it from many other indole-based or receptor-limited molecules [72]. The observed inverse alignment between intracellular melatonin accumulation and ROS levels suggests that melatonin’s antioxidant activity may result from this direct cytosolic presence, allowing it to neutralize free radicals before they inflict oxidative damage [73]. Disruption of redox balance, particularly elevated ROS levels, can precipitate anoikis, a form of programmed cell death triggered by detachment from the extracellular matrix [74]. Thus, melatonin’s modulation of intracellular ROS may influence anoikis susceptibility in colorectal cancer cells.

Despite the multidisciplinary approach used to elucidate melatonin’s anticancer mechanisms, this study encounters limitations. Most analyses were performed at the RNA level, which constrains inferences about protein abundance and function. Future work should incorporate proteomics and targeted protein assays (e.g., immunoblotting, FRET) to determine whether the observed transcriptomic changes are reflected at the protein level. In addition, employing advanced methodologies, such as CRISPR-mediated receptor knockouts and TF occupancy would help validate causal mechanisms and deepen our understanding of melatonin’s antitumor actions.

## 5. Conclusion

Melatonin represses a defined subset of the mitotic programme that is upregulated across all stages of CRC, lowering *BUB1* and *CCNA2* while leaving *NCAPG* unchanged, the pattern predicted if NF-Y- dependent transcription is reduced, and the reason *NCAPG* functions here as an internal specificity control rather than a negative result. The associated G1 accumulation, S-phase depletion and moderate apoptosis are consistent with loss of cyclin A2. The NF-Y model remains a hypothesis until occupancy and subunit-level data are available. Independently of it, β2-microglobulin is unsuitable as a reference gene in melatonin-treated CRC cells, and its induction is a specific, testable lead.

## 6. List of abbreviations

CRC: Colorectal Cancer
TNM: Tumour-node-metastasis
GPCR: G protein-coupled receptor
qPCR: Quantitative Polymerase Chain Reaction
MTR1: Melatonin receptor
CTCF: Corrected total cell fluorescence
MTR2: Melatonin receptor 2
IC: Inhibitory Concentration
cAMP: Cyclic Adenosine Monophosphate
PKA: Protein kinase A ERK Extracellular signal-regulated kinase
TF: Transcription factor
DEG: Differentially expressed genes
PPI: Protein-Protein Interaction
GEO: Gene Expression Omnibus
LCM: Laser Capture Microdissection
STRING: Search Tool for the Retrieval of Interacting Genes
MCC: Maximal Clique Centrality
MCODE: Molecular Complex Detection
KEGG: Kyoto Encyclopaedia of Genes and Genomes
miRNA: MicroRNA
GO: Gene Ontology
GSEA: Gene Set Enrichment Analysis
X2K: Expression2Kinases
GEO2R: Gene Expression Omnibus 2R
IHC: Immunohistochemistry
HPA: Human Protein Atlas
DMEM: Dulbecco’s Modified Eagle Medium
FBS: Fetal Bovine Serum
PBS: Phosphate-Buffered Saline
DMSO: Dimethyl Sulfoxide
cDNA: Complementary DNA
Ct: Threshold cycle
PI: Propidium Iodide
AF3: AlphaFold 3
TPM: Transcripts Per Million
COAD: Colon Adenocarcinoma
Ex/Em: Excitation/Emission
DCF: 2′,7′-Dichlorofluorescein
CDK2: Cyclin dependent kinase 2
PepT2: Peptide Transporter 2
NP: Nanoparticle
DSI: Disease Specificity Index
DPI: Disease Pleiotropy Index
RPU: Relative Pixel Units
MFI: Mean Fluorescence Intensity
TYMS: Thymidylate synthase

## 7. Declarations

### 7.1 Supplementary Materials

Supplementary Material provided along with the manuscript and cited wherever related.

### 7.2 Authors’ Contributions

FR has done Conceptualization, Experimental Design, Investigation, Writing–Original Draft, Figures and Tables Preparation. BD performed Experimental Design, Methodology, and Supervision. KK contributed to Providing Resources, Supervision and Administration. All authors read and approved the final manuscript.

### 7.3 Funding

Not applicable

### 7.4 Ethics Declaration

Not applicable

### 7.5 Consent for Publication

Not applicable

### 7.6 Availability of Data

All data used and analysed in this paper are freely available in the mentioned databases e.g. Gene Expression Omnibus (https://www.ncbi.nlm.nih.gov/geo/). **Figure 6C** and **Figure 7** are self-generated illustrations designed by BioRender (https://www.biorender.com/).

## Acknowledgement

We gratefully thank National Institute for Medical Research and Development (NIMAD, project 962487) and research council of Tarbiat Modares University for their administrative assistance. The research has been supported by Research Organization of Tarbiat Modares University as part of the support provided for MSc thesis projects.

## 7.7 Conflict of Interests

All authors declare that they have no conflict of interests.

## 7.8 Additional Information

This research has been conducted as the thesis project of the first MSc degree of the first author (FR), in partial fulfilment of the requirements for degree Master of Science (MSc) in Biochemistry at the Faculty of Biological Sciences, Tarbiat Modares University, Tehran, Iran. Contact lead.

## Notes

### Competing Interest Statement

The authors have declared no competing interest.

