## Supplementary 1 for "Deciphering Novel Transcriptional Wiring in Colorectal Cancer: An Integrative Bioinformatic and Experimental Study"

$$\textbf{Cell Viability (\%)} = \frac{\left(\text{Mean Abs}^{\text{treated}}_{570\,\text{nm}} - \text{Mean Abs}^{\text{treated}}_{630\,\text{nm}}\right) \times 100}{\left(\text{Mean Abs}^{\text{control}}_{570\,\text{nm}} - \text{Mean Abs}^{\text{control}}_{630\,\text{nm}}\right)}$$

**Supplementary 1A.** Cell Viability Calculation Based on MTT Absorbance Readings [1].

---

$$\textbf{Cell Viability (\%)} = \frac{\text{Top} - \text{Bottom}}{1 + 10^{((\log_{10} \text{IC}_{50} - \log_{10}[\text{Melatonin}]) \times \text{Hill Slope})}}$$

**Supplementary 1B.** Dose-Response Curve (Four-Parameter Logistic Model).

---

$$\textbf{Relative Expression Ratio} = (E_{\text{target}})^{\Delta C_t\ (\text{control}-\text{sample})} \Big/ (E_{\text{ref}})^{\Delta C_t\ (\text{control}-\text{sample})}$$

Where:

- $E_{\text{target}}$ is the amplification efficiency of the target gene
- $E_{\text{ref}}$ is the amplification efficiency of the reference gene
- $\Delta C_t$ represents the difference in threshold cycles between control and treated samples for each gene.

**Supplementary 1C**. Relative Gene Expression Quantification Using the Pfaffl Method

---

$$\mathbf{CTCF} = \text{Integrated Density} - (\text{Area of selected cell} \times \text{Mean fluorescence of background})$$

Where:

- **Integrated Density** refers to the sum of the pixel values within the selected cell region
- **Area of selected cell** is measured in pixels
- **Mean fluorescence of background** is calculated from regions lacking cells to account for non-specific signal

**Supplementary 1D.** Corrected Total Cell Fluorescence (CTCF) Calculation

---

### Refs:


1. Mosmann, T., *Rapid colorimetric assay for cellular growth and survival: application to proliferation and cytotoxicity assays*. Journal of immunological methods, 1983. **65**(1-2): p. 55-63.
